# The Brain, Body, and Behavior Dataset (BBBD): Multimodal Recordings during Educational Videos

**DOI:** 10.1101/2025.04.29.651259

**Authors:** Jens Madsen, Nikhil Kuppa, Lucas C. Parra

## Abstract

When humans engage with video, their brain and body interact in response to sensory input. To investigate these interactions, we recorded and are releasing a dataset from N=178 participants across five experiments featuring short online educational videos. This dataset comprises approximately 110 hours of multimodal data including electrocardiogram (ECG), heart rate, respiration, breathing rate, pupil size, electrooculogram (EOG), gaze position, saccades, blinks, fixations, head movement, and electroencephalogram (EEG). Participants viewed 3-6 videos (mean total duration: 28±5 min) to test attentional states (attentive vs. distracted), memory retention (multiple-choice questions), learning scenarios (incidental vs. intentional), and an intervention (monetary incentive). Demographic data, ADHD self-report (ASRS), and working memory assessments (digit span) were collected. Basic statistics and noteworthy effects: increased alpha power in a distracted condition, broadband EEG power increases from posterior to anterior scalp, increased blink-rate, and decreased saccade-rate in distracted and intervention conditions. All modalities are time-aligned with stimuli and standardized using BIDS, making the dataset valuable for researchers investigating attention, memory, and learning in naturalistic settings.

## Background & Summary

The increasing availability of large, high-quality datasets, driven by the principles of open science^1^ has catalyzed interdisciplinary research. Neuroimaging provides detailed insights into brain activity^2^, while physiological studies reveal dynamic bodily responses to stimuli^3,4^. Research into brain-body interactions bridges these domains, linking neural processes with physiological states^5^. This work is complemented by efforts to link stimulus properties with neural^6,7^ and physiological responses^8,9^. Naturalistic experimental designs^10^ have enhanced the potential for cross-disciplinary research. By leveraging multimodal datasets within reproducible frameworks, researchers have decoded complex cognitive processes involving attention^11^, memory^12,13^, and learning^14^.

Our lab takes a distinct approach by integrating neural, behavioral, and physiological modalities with precise time alignment across modalities and with the stimulus. This enables detailed modeling of interactions between modalities and their relationship to the stimulus, surpassing the limitations of single-technique datasets. For instance, we demonstrated that synchronized eye movements predict test performance when students watch educational videos^15^. Additionally, we showed that audiovisual narratives synchronize heart rate, with modulation by attention and conscious narrative processing^16^. By integrating modalities, we revealed that intersubject correlation (ISC) of physiological, neural, and behavioral signals is interconnected^17^. High ISC in EEG corresponds to high ISC in heart rate, underscoring the dynamic interplay between brain and body.

While the rise in multimodal datasets using naturalistic stimuli is encouraging, there remains a notable gap in datasets within the audiovisual domain. Several datasets focus on movie watching while collecting single modalities in addition to fMRI, e.g. pulse oximetry (N=20)^18^ or eye tracking (N=15)^19^ while watching the feature film *Forrest Gump*. Other datasets were recorded under the exposure of different participants to the same movie using different modalities (fMRI and MEG, N=11)^20^. In other studies participants (N=51) watched audiovisual stimuli while their intracranial electroencephalography (iEEG) and fMRI were acquired in repeated presentations^21^. Lastly, in auditory experiments multiple neural measures have been used e.g. recording fMRI and MEG while subjects listened to an audio in repeated presentations^20^ (N=12).

A few datasets include both neural and physiological measures. For instance, the K-EmoCon dataset combines EEG with peripheral physiological signals^21^ (N=32), offering the advantage of data collected in uncontrolled, real-world environments. However, the lack of common stimuli or triggers limits the potential for cross-subject analyses. The DEAP dataset (Dataset for Emotion Analysis using Physiological Signals)^24^ focuses on affective responses to short music video clips (N=32), recording a range of physiological and neural signals, including electrodermal activity (EDA), heart rate (HR), skin temperature, electromyography (EMG), electrooculogram (EOG), EEG, respiration, and frontal-facing video.

Despite their merits, none of these datasets integrate all the following elements: audiovisual stimuli, neuroimaging, physiological and behavioral data, and memory assessment information. Such a combination enables a comprehensive analysis of how the brain and body interact in response to naturalistic audiovisual stimuli.

Our dataset bridges these gaps. BBBD offers a comprehensive and time-aligned collection of physiological, behavioral and neural measures, alongside responses to questionnaires and detailed demographics metadata. Physiological measures in the dataset include electrocardiogram (ECG), EOG, respiration, heart rate, breathing rate, heartbeat timestamps, breath peak (inspiration) timestamps, and pupil size; behavioral measures include gaze position, saccades and saccade rate, blinks and blink rate, gaze fixations and fixation rate, and head position (horizontal, vertical, and depth); and neural activity was recorded using EEG.

Beyond the breadth of physiological and behavioral data, our dataset incorporates key experimental features: memory questions tailored to each story viewed, attentional manipulation conditions (attentive vs. distracted viewing) and participant characterization through demographics, ADHD self-report scale (ASRS) scores evaluation^25^, and working memory scores assessed using the Digit Span memory task^26^.

This dataset’s vast majority, approximately 97% of continuous signal data, is being newly published in a standardized data structure following the Brain Imaging Data Structure’s (BIDS) convention. A limited subset of this dataset that was released in its preliminary form (Table 1), featured in three prior studies conducted by our group^15–17^.

**Table 1:**
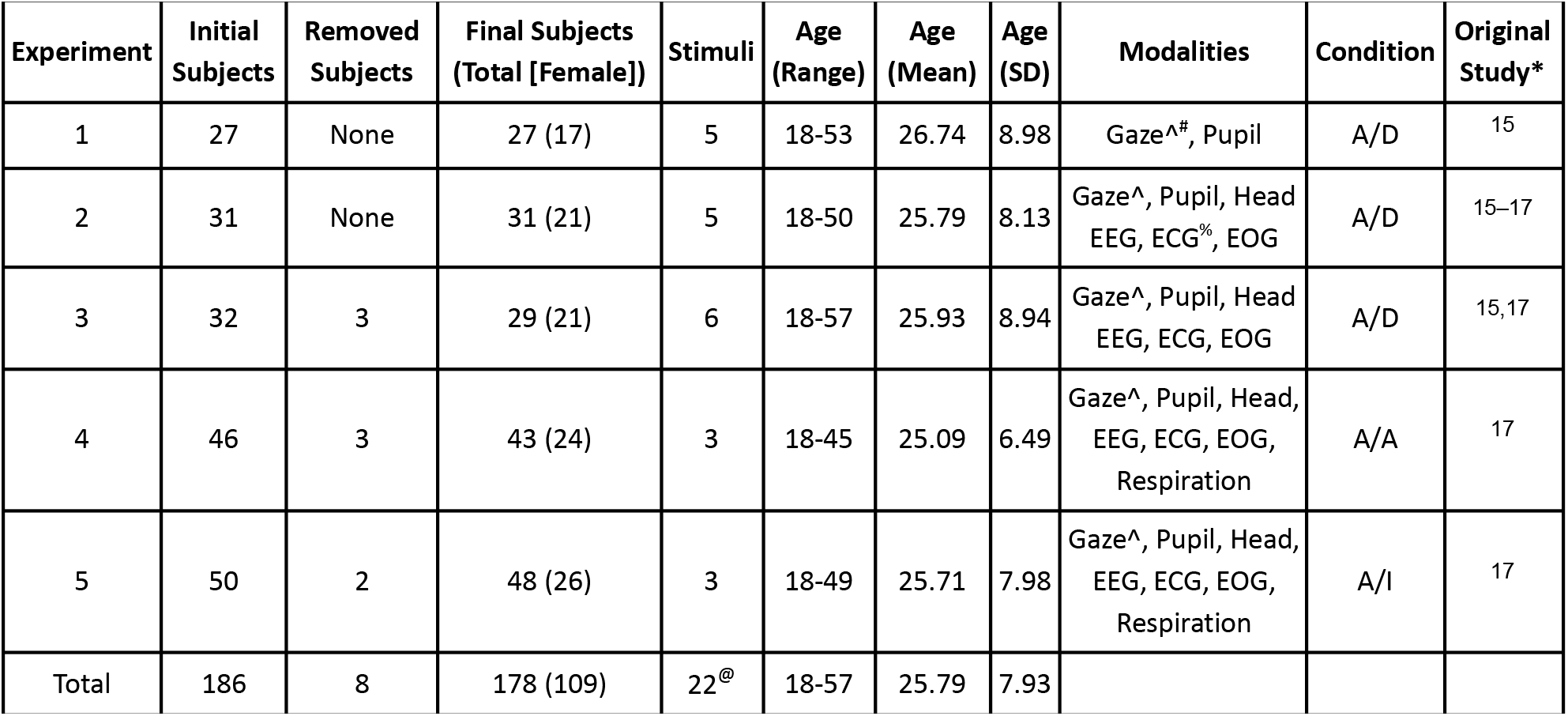
Dataset summary. * Original studies may have reported on a different number of subjects from what is being released here. ^ includes blinks, saccades, fixations, saccare rate, blink rate. In the condition column, **A** indicates the dataset contains the Attend condition where subjects watched the videos normally, **D** indicates the Distract condition, where the subject carried out an arithmetic task silently in their heads and **I** is the Intervention condition where participants were told they would earn reward for answering questions correctly about the video. The current data is newly published with the exception of (#) raw gaze position and pupil data for 2 of 10 sessions (2 videos in the attend condition) in Experiment 1^15^ and (%) ECG data for Experiment 2^16^. (**@**) Among the 22 total stimuli presented across 5 experiments, 11 stimuli were unique.

## Methods

### Participants

A total of 180 subjects participated across 5 experiments, where they viewed short explanation videos. Participants were recruited through craigslist.org’s New York City region, as well as from the mailing lists of the student body of the City College of New York (CCNY) for these experiments. CCNY has a diverse student body in terms of age, ethnicity, primary languages, skills, etc., and participants from the public in New York City only added to this diversity. Subjects were compensated for their time at a rate of $20 USD per hour, with the experiment lasting between 1 and 2 h. Table 1 summarises the participant information experiment wise, and also lists the modalities that were collected under respective conditions.

### Stimuli

Across 5 experiments, 11 unique video stimuli were shown to the subjects (Table 2) These stimuli have been selected from YouTube channels that post short explanation videos on a number of educational topics and have been used in previous studies^15–17^.

**Table 2:** Explanation videos used, education channels they were sourced from, duration, topic name, and their respective YouTube video IDs. To access these videos on YouTube, attach a video ID of interest to this link https://www.youtube.com/watch?v=

| Stimuli name | Experiment | Channel name | Duration | Topics | Youtube ID |
| --- | --- | --- | --- | --- | --- |
| Why are Stars Star-Shaped | 1,2,4,5 | minutephysics | 3:29 | Physics | VVAKFJ8VVp4 |
| How Modern Light Bulbs Work | 1,2,4 | minutephysics | 2:23 | Physics | oCEKMEeZXug |
| The Immune System Explained – Bacteria | 1,2,4,5 | Kurzgesagt - In a Nutshell | 7:49 | Biology | zQGOcOUBi6s |
| Who Invented the Internet - And Why | 1,2 | Kurzgesagt - In a Nutshell | 6:33 | Computer Science | 21eFwbb48sE |
| Why Do We Have More Boys Than Girls | 1,2 | MinuteEarth | 2:34 | Biology | 3IaYhG11ckA |
| What If We Killed All the Mosquitoes | 3,4 | What If | 4:47 | Biology | 9w-5wJYVmcw |
| Are We All Related | 3,4,5 | Be Smart | 6:26 | Biology | mnYSMhR3jCI |
| Work and the Work-Energy Principle | 3 | Khan academy | 5:48 | Physics | 30o4omX5qfo |
| Dielectrics in Capacitors Circuits | 3 | Khan Academy | 6:27 | Physics | rkntp3_cZl4 |
| How Do People Measure Planets & Suns | 3 | eHowEducation | 4:39 | Astronomy | bYgV9nvgJ3E |
| Three Factors That May Alter the Action of an Enzyme | 3,4 | eHowEducation | 3:12 | Biology | IkRZKqDdwzU |

### Procedure

All experiments were carried out at the City College of New York with the required ethical approval from the Institutional Review Boards (IRB) of the City University of New York. Informed consent was obtained from all participants prior to the experiment. The resulting anonymized dataset has been made publicly available in accordance with these ethical and institutional guidelines. Subjects were seated comfortably in a sound-attenuated booth with white fabric walls and normal ambient LED lighting around, and all data acquisition devices were securely and safely attached to participants. They watched the videos on a 27” monitor approximately 60 cm from the subject, while audio was delivered through stereo speakers placed next to the monitor and separated by 60° from the subject (Fig. 1).

**Figure 1:**
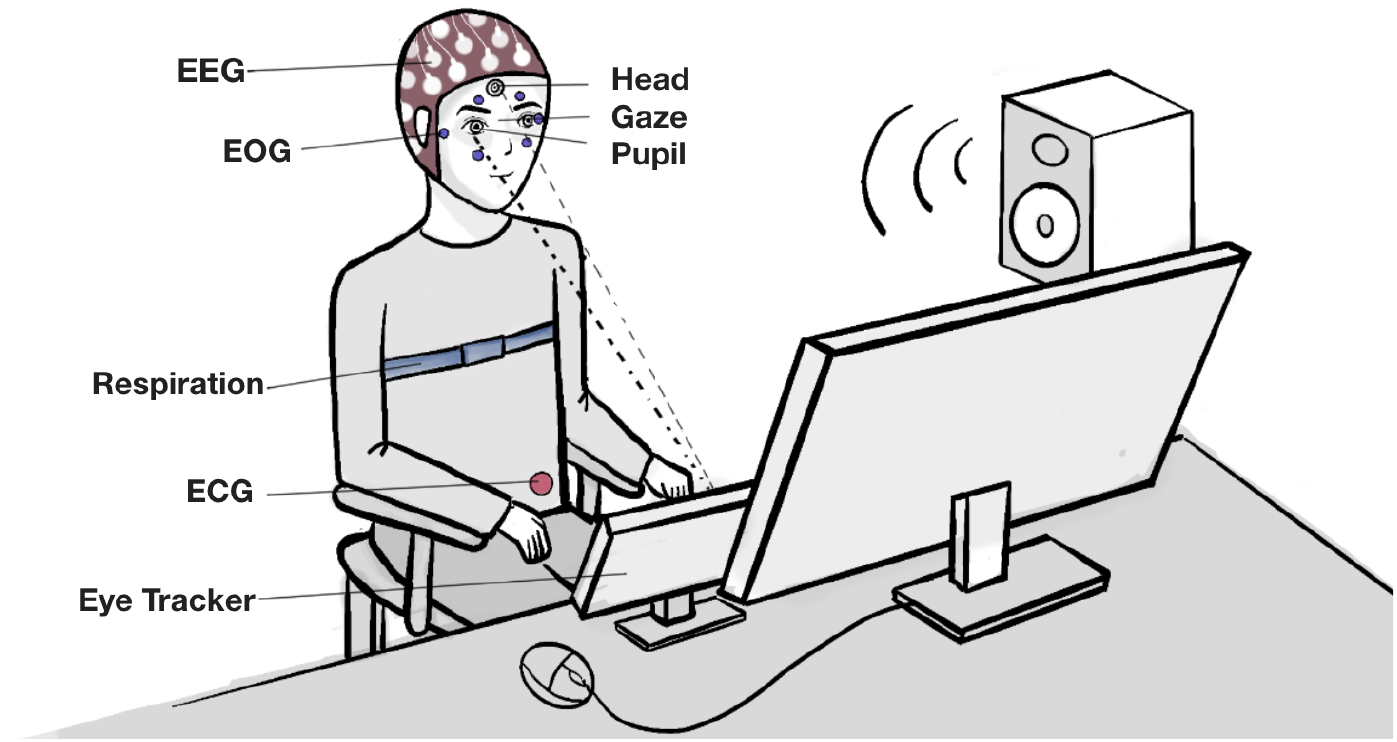
Illustrated figure of the experiment setup. The pupil, gaze and head position data is collected by the eye tracker placed at the bottom of the monitor, respiration data is collected by the belt worn on the chest, and the EEG data is collected by the 64-channel cap. EOG and ECG are collected from their respective BioSemi ActiveTwo electrodes.

In all experiments, subjects were instructed to watch the videos normally as they would at home, while being relaxed and sitting still. We refer to this as the attentive condition. In the distracted condition, subjects were asked to watch the videos again, except this time they were assigned the Descending Subtraction Task^27^ with slight variation: count backwards in their minds from a prime number between 800 and 1,000 chosen randomly, in steps of 7. This task was incorporated with a goal of distracting subjects from the stimulus without requiring noticeable responses. Serial subtraction tasks such as this are typically used to assess mental capacity and have also been used to assess attention.

The attentive condition was followed by a quiz (10-12 questions per video) pertaining to the factual information that was imparted in the video. Within an experiment, the presentation order of all videos and questions were randomized across subjects to minimize order effect^28^. In addition to the questions regarding the content of the videos, subjects were also asked 3 engagement questions to understand how they perceived each video and whether they found it informative.

### Experimental procedures

Experiments 1-3 were designed to assess either intentional learning (subjects were informed in advance that they would be tested on the contents of the video) or incidental learning (subjects were not made aware they would be tested on the contents of the video). The quizzes for each video are short with 10-12 four-alternative forced-choice questions. For the Intentional Learning condition, a quiz is followed after each video, whereas in the Incidental Learning condition, a cumulative quiz is given after all the videos have been watched (Fig. 2).

**Figure 2:**
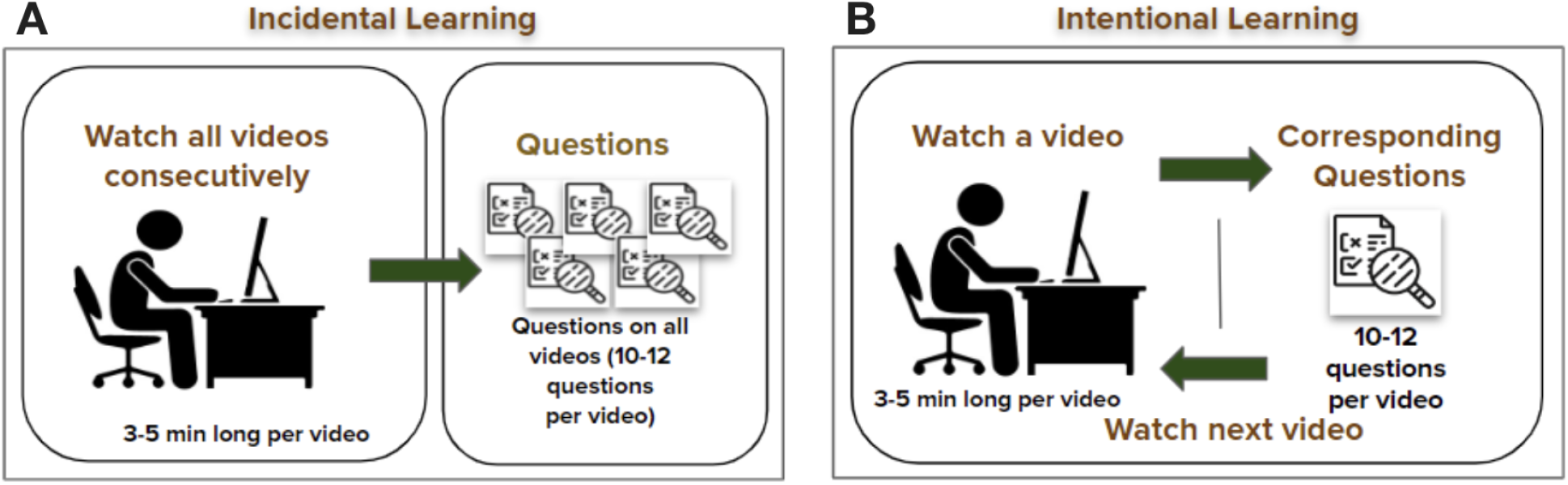
Experiments 1-3 with two learning conditions: **A)** <u>Incidental Learning</u>: Subjects watch all videos consecutively, and then take a quiz pertaining to the videos. Subjects *did NOT know* they would be tested on the video content. **B)** <u>Intentional Learning</u>: Subject watches one video at a time, answers questions pertaining to the video and does the same for other videos. Subjects *did know* they would be tested

Experiments 4 and 5 were designed to have a slight variation in the incidental and intentional learning respectively. Experiment 5 in particular had an intervention between the two sessions (Fig 3).

**Figure 3:**
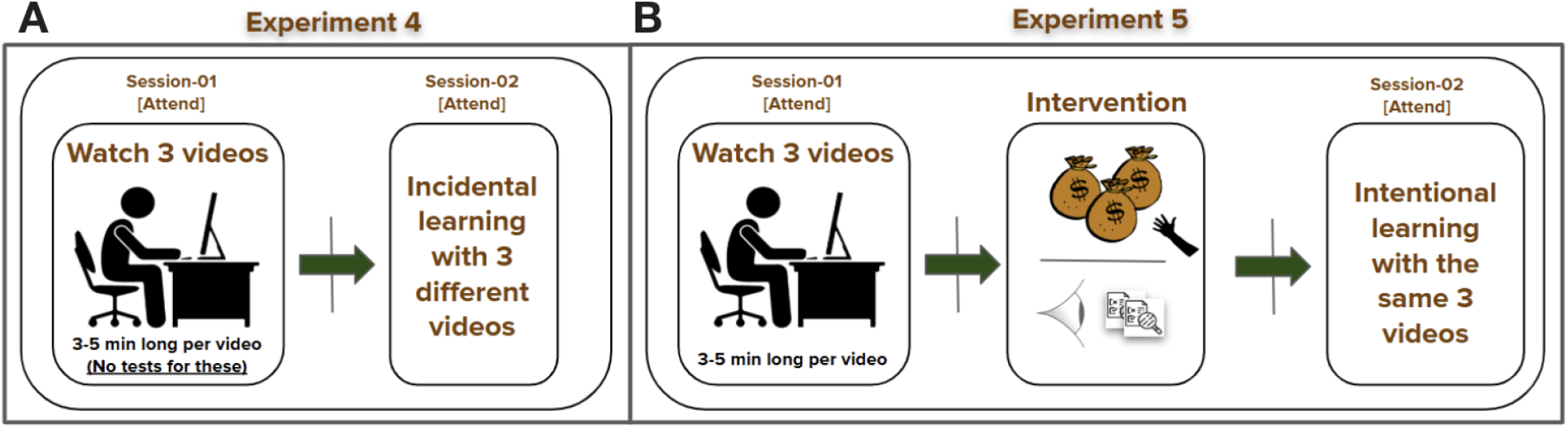
**(A)** Experiment 4 and **(B)** Experiment 5. During Experiment 5, subjects were randomized to an intervention or control condition. In the intervention condition, they receive monetary incentive for good, correctly answering questions about the video, and questions were shown prior to the video so they knew what to attend to.

#### Experiment 1

In Experiment 1 subjects were informed they would be questioned about the contents of the video (**intentional learning)**. After they watched all the videos, they watched them again in a distracted condition. Subjects watched 5 different informative videos while tracking their gaze and pupil size. The position of gaze is an overt indicator of attention and this information helps researchers understand what points of the video they’re focused on and at what points their mind starts wandering away.

#### Experiment 2

Experiment 2 was **incidental learning** - where the subjects watched all the videos not knowing they would be questioned about the content of each video all together at the end of the video-watching. However, subjects were also given a questionnaire before being exposed to stimuli to test their domain knowledge pertaining to the video topics. After they had watched the videos and answered the questions they watched the videos again in the distracted condition.

For this experiment we collected physiological data - ECG, EEG, Head motion, along with gaze and pupil size. Subjects watched 5 informative stimuli.

#### Experiment 3

In experiment 3, subjects were aware they would be tested on the material of each video presented (**Intentional Learning**). We tested 6 informational video stimuli, 2 each from 3 different styles of presentation for this experiment to understand if the video design would have an impact on the attention retention of students. The video styles tested were

1. Presenter and animation
2. Presenter and glassboard
3. Animation and writing hand.

After they watched the 6 videos in the attentive condition they watched them again in the distracted conditions. We record the same data from the subjects as in experiment 2.

#### Experiment 4

For experiment 4, subjects watched 6 videos in an attentive condition. They were tested on the last 3 videos they watched. Subjects were not informed they would be tested on the content, constituting this incidental learning. During the experiment we recorded EEG, EOG, ECG, respiration, pupillary responses, gaze, and head position throughout.

#### Experiment 5

Finally, for experiment 5, subjects were instructed to watch 3 instructional videos in an attentive condition twice, while their EEG, EOG, ECG, respiration, pupillary responses, gaze, and head position were recorded. Between the two conditions there was an intervention - subjects were incentivised with money for every question they got right at the test that was given to them at the end of the second condition. They were also shown what questions were going to be asked, and were made to watch the same videos again.

The first attentive condition can be interpreted as incidental learning as subjects were not told they would be asked questions, and the second attentive condition can be interpreted as a slight deviation from intentional learning since they were told they would be tested, and were shown what they’d be asked. We track the same signals as in experiment 4.

##### Note for Exp 4 & 5

Some EEG signals in this experiment have a completely electrical component of 16 Hz, and is not a physiological artifact. This issue arose during the experimental setup due to unforeseen electrical noise in the Biosemi system.

### Signal Acquisition

While signals were recorded by different devices at different sampling rates, for the final data release, all signals have been down-sampled to 128 Hz and are time-synchronized.

#### Recording and preprocessing of EEG

EEG data was acquired at 2,048 Hz using the BioSemi ActiveTwo system with a 64-electrode cap (10/10 system) with ground electrodes near POz. Signals are band-pass filtered (0.016–25 Hz) by the analog device before sampling, then digitally high-pass filtered (0.05 Hz) and notch-filtered at 60 Hz to remove line noise. Robust PCA^29^ was used for artifact removal, followed by low-pass filtering (64 Hz) and down-sampling to 128 Hz. On average across all EEG channels and sessions, in Experiment 2 approximately 0.84% (SD = 2.62%) of the raw EEG was marked as artifacts and corrected, 1.75% (SD = 6.09%) was corrected in Experiment 3, 0.32% (SD = 0.84%) was corrected in Experiment 4, and 0.62% (SD = 1.83%) was corrected in Experiment 5.

EOG artifacts were subtracted using least-squares noise cancellation via linear regression of EOG signals. Outliers (values exceeding four times the interquartile range of the median-centered signal) were replaced with interpolated samples including samples within a 40 ms window using neighboring electrodes^17^.

#### Recording and preprocessing of EOG

EOG was recorded with 6 auxiliary electrodes (located above, below, and laterally to each eye). However, the exact mapping of these electrode numbers to their corresponding locations is inconsistent across participants. This data was pre processed the same way as EEG^17^.

#### Recording and preprocessing of ECG

ECG was recorded at 2,048 Hz using the same BioSemi ActiveTwo system with electrodes below the left collarbone and on the left lumbar region. Analog filtering is the same as EEG. Digital signals were high-pass filtered (0.5 Hz cutoff) and notch-filtered at 60 Hz to remove line noise. R-wave peaks were identified using MATLAB’s findpeaks function, and manually corrected using a graphical interface (all timestamps are included in this release). The time stamps for the R-peaks represent times relative to the start of the video stimulus. However, these timestamps were computed based on the filtered signal and using the source sampling frequency of 2,048Hz and may not align precisely with the apparent peak of the downsampled signals at 128Hz. Instantaneous heart rate (HR) was calculated for each beat as the inverse of intervals between successive R-peaks. The HR signal was resampled to 128 Hz for uniformity across subjects^17^.

#### Recording and preprocessing of gaze position, head position, and pupil size

Gaze position, head movements, and pupil size were recorded at 500 Hz using the Eyelink 1000 eye tracker (SR Research Ltd.) with a 35mm lens. Subjects were seated comfortably without a chin rest, and a standard 9-point calibration scheme with manual verification was used. Calibration and instruction screens matched the average luminance of the experiment videos to ensure stable pupillary responses. Drift checks were performed after each condition (attend, distract and intervention) with recalibration required if angular error exceeded 2°.

Blinks and saccades were detected using the eye tracker’s algorithm that is used when saving to BDF files (the algorithms used during streaming via Ethernet are less precise). Additional artifacts were detected by finding outliers (4 times the IQR) from a detrended pupil signal (using a 200ms median filter). Blinks, artifacts, and the 100 ms before and after these events, were corrected with linear interpolation. On average, about 22% (SD = 19%) of every gaze file and about 32% (SD = 25%) of every pupil file were identified as artifacts and corrected accordingly (N = 1455 files for both gaze and pupil).

Instantaneous saccade rate, calculated for each saccade as the inverse of intervals between saccades, was resampled to 500 Hz. Head position was tracked using a forehead sticker (Fig. 1), with X and Y coordinates representing target position in camera sensor pixels and Z indicating distance from the tracker in millimeters. All signals were resampled to 128 Hz for consistency^17^.

#### Recording and preprocessing of respiration

Respiration signal was recorded at 2,048 Hz using the BioSemi ActiveTwo system and SleepSense 1387 Respiratory Effort Sensors, with a belt worn around the chest to measure tension. Signal polarity was corrected by detecting peaks and inverting the phase as needed. Signals were resampled to 128 Hz^17^.

#### Synchronizing signals across modalities

Common onset and offset triggers were used for the segmentation of the physiological and neural signals, recorded with a BioSemi ActiveTwo system. A flash and beep sounds were embedded at the beginning and end of each stimulus to validate precise alignment of the digital triggers across all subjects, and these sounds were captured using a StimTracker (Cedrus) which is recorded as digital triggers with the BioSemi system. In addition to the BioSemi recording system, the triggers were also sent to the eye tracking recording system. The timestamps recorded in each of these systems were used to harmonize onset and duration with the common triggers (by assuming a linear transformation of times). Some of the data was recorded directly to a BDF file through the software provided by the manufacturer, while others used the Lab streaming layer (LSL) protocol.

## Data Records

### Data access

All data can be directly accessed through a dedicated website for this dataset (https://bbbd.pythonanywhere.com/). This website provides links to directly download all the neural, physiological and behavioral data - hosted on the International Neuroimaging Data-Sharing Initiative (INDI)’s Amazon Simple Storage Service (S3) infrastructure. INDI works in collaboration with the Neuroimaging Tools & Resources Collaboratory (NITRC), and ensures long-term accessibility and compliance with community data-sharing standards. BBBD is officially listed in the INDI dataset repository (see https://fcon_1000.projects.nitrc.org/indi/IndiRetro.html) under *“The Brain, Body, and Behavior Dataset (BBBD): Multimodal Recordings during Educational Videos*.*”*

We included the list of stimuli used for each experiment along with their Youtube URLs in Table 1, and also within the respective README files located in the root directories of each experiment (can be accessed from the BIDS-structure download links on the website). For every experiment, individual .mat files are also provided for each modality - both raw and preprocessed data. Each of these files comes with a README variable containing all the information necessary to understand and work with the data.

### Data organization

To make this dataset easily accessible and comprehensible, we reformatted it into the BIDS data structure^30^. Each of the five datasets in the BIDS format has files that describe the dataset, its participants, related metadata - dataset_description.json, participants.tsv and participants.json, and questionnaires (quizzes, Adult ADHD Self Report (ASRS) test, digit span scores, video engagement) providing essential information about the study and participants (located in the /phenotype directory). Each participant’s data is organized by subjects and then further divided into sessions to accommodate multi-session data collection.

The standard dictates that any non-raw data is put into a derivatives directory, this contains processed data derived from the raw recordings, continuous data such as heart rate, saccade rate or discrete data like saccades and blinks along with preprocessed and cleaned physiological signals. Inside each session folder, one may find modality-specific subfolders, which contain data as mentioned in Table 3.

**Table 3:** Data organization in the subdirectories.

| Derived (/derivatives) Subdirectories |  |  | Raw (Root Directory) Subdirectories |  |  |
| --- | --- | --- | --- | --- | --- |
| /eeg | /beh | /eyetrack | /eeg | /beh | /eyetrack |
| Pre-processed EEG | Pre-processed ECG | Pre-processed Pupil-size, Gaze | EEG | ECG | Gaze Coordinates |
|  | Heart-rate,<br>Breath-rate<br>(continuous signal) | Fixation-rate, Saccade-rate, Blink-rate<br>(continuous signal) | Channels, Electrodes,<br>Coordinates,<br>Event logs | EOG | Pupil Size |
|  | R-Peaks,<br>Breath Peaks<br>(timestamps) | Fixations<br>(start and end timestamps,<br>visual degree angles in x and y) | Metadata (.json) | Respiration | Head Position |
|  |  | Saccades, Blinks<br>(start and end timestamps) |  | Metadata (.json) | Metadata (.json) |

The figure below is a detailed breakdown on the total minutes of raw data available for each session across every experiment (Fig 4).

**Figure 4:**
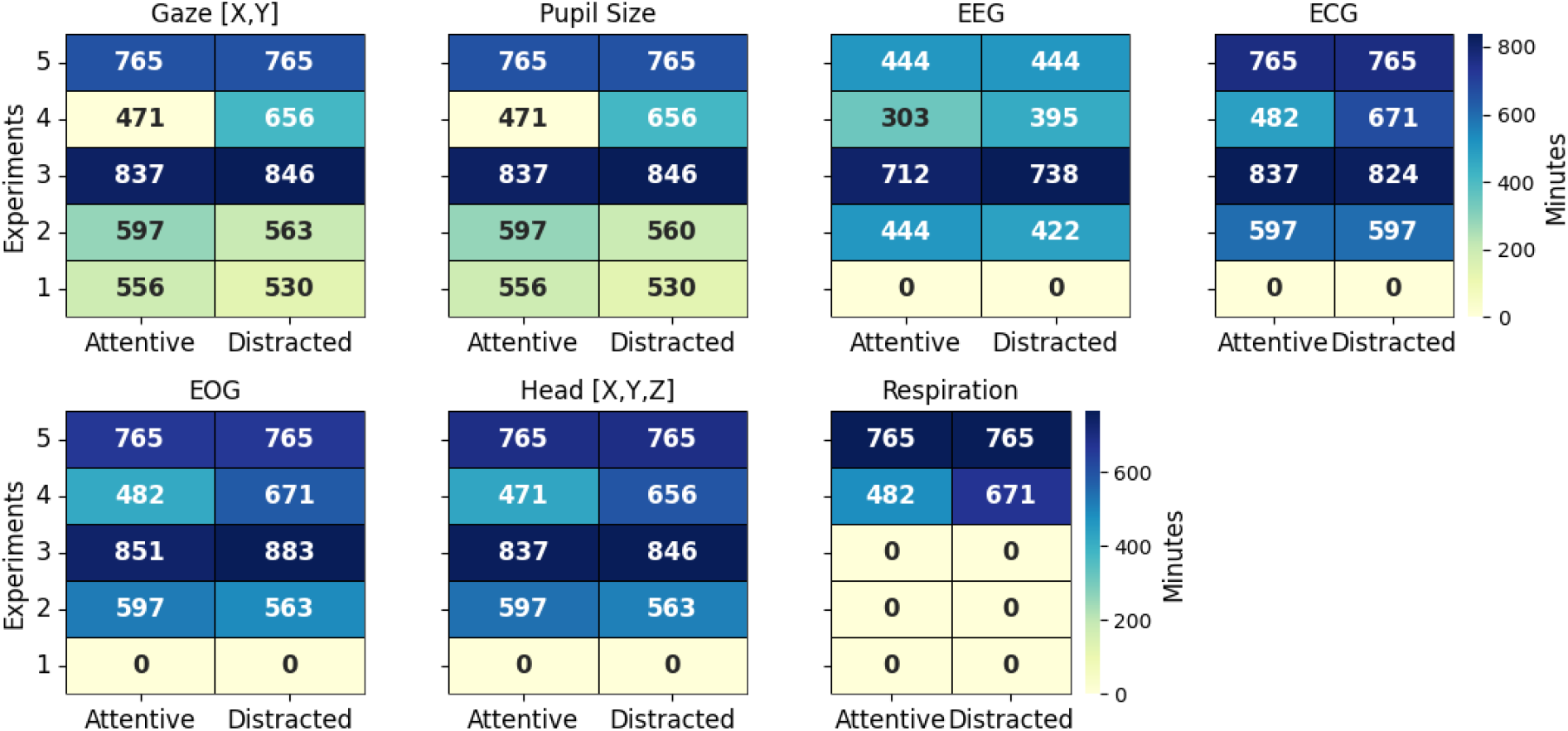
Duration of available raw data per session (Attentive and Distracted) across all 5 experiments for each modality (in minutes) <u>Note</u>: Non-EEG **raw data** files are stored in a compressed tsv format (**tsv.gz**), and these files do not contain a header due to metadata availability. Non-EEG **derived data** files are stored in a **tsv** format without compression, and they do have a header.

## Technical Validation

All statistical results presented in this section (Heart Rate, Heart Rate Variability, Blink Rate, Saccade Rate, Pupil Size) reflect within-subject paired t-tests conducted independently for each experiment.

### EEG

To validate the quality of the EEG data we computed the power spectral density averaged across subjects and conditions for each experiment (Fig. 5A). We observed the expected 1/f spectrum with a peak in the alpha band (8-12 Hz). We note a gradual decrease in power across all frequencies as we go from frontal to occipital electrodes. During rest, this phenomenon has previously been observed at frequencies below the alpha band^31^. Here it is consistently observed during video watching across all bands up to 20 Hz.

**Figure 5:**
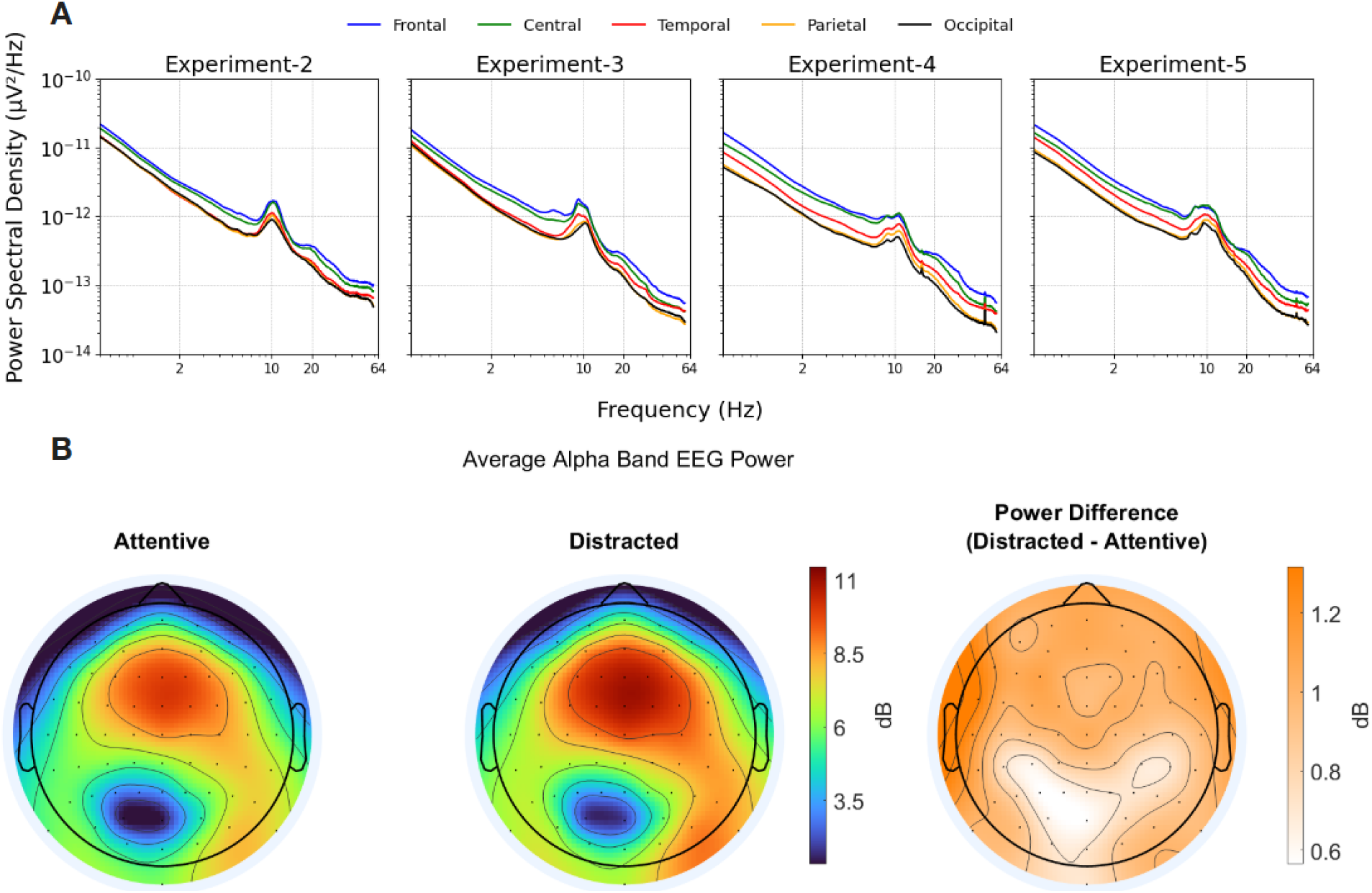
**(A)** Power Spectral Density (log-log) across brain regions for all experiments with the preprocessed EEG data averaged over each region across all participants in the experiments. **(B)** Scalp distribution of the average power (in dB) across the alpha band (8-12 Hz) of the 64 channels of electroencephalograph (EEG) recorded on all subjects in Experiments 2 and 3, for attentive and distracted conditions respectively. The difference between the powers of the two sessions is also illustrated.

We also observed that the alpha band power is higher in the Distracted condition compared to the Attending condition across the entire scalp (Fig 5B). This is expected, because directing attention away from an external stimulus is well-known to increase alpha power. ^32^

### ECG, R-Wave Peaks, Heart Rate

ECG signals, the detected R-Wave peaks along with the corresponding heartbeats were visually checked for any artifacts. Artifacts in the heart rates mostly originated due to incorrectly detected R-Peaks, and these peaks were corrected manually, thereby fixing the heart rate. Nonetheless, for some subjects and conditions there are residual artefacts in the HR signal due to missing data or movement artefacts in the ECG. Heart rate (HR) did not differ significantly between conditions for Experiments 2 and 3, but was elevated in the intervention condition for experiment 4 (Fig. 6A, t(28)=-0.37, p=0.72, d’=-0.07; t(27)=-1.74, p=0.093, d’=-0.34; t(47)=4.05, p=1.9·10^−4^, d’=0.59, within-subject paired t-test). Distracted conditions in Experiments 2 and 3 had no significant impact on the Heart rate variability (HRV), but the intervention condition in Experiment 5 showed a statistically significant decrease in the HRV (Fig. 6B, t(28)=0.07, p=0.95, d’=0.01; t(27)=1.68, p=0.105, d’=0.32; t(47)=-4.38, p=6.6·10^−5^, d’=-0.64, within-subject paired t-test). Previous studies show that students have a reduction in their HRV when they take a test^33^, and our results seem to follow a similar pattern where subjects, in anticipation of excelling the tests to gain monetary reward, ended up having a lower HRV. HRV has been shown to increase for people with higher Working Memory Capacity^34^.

**Figure 6:**
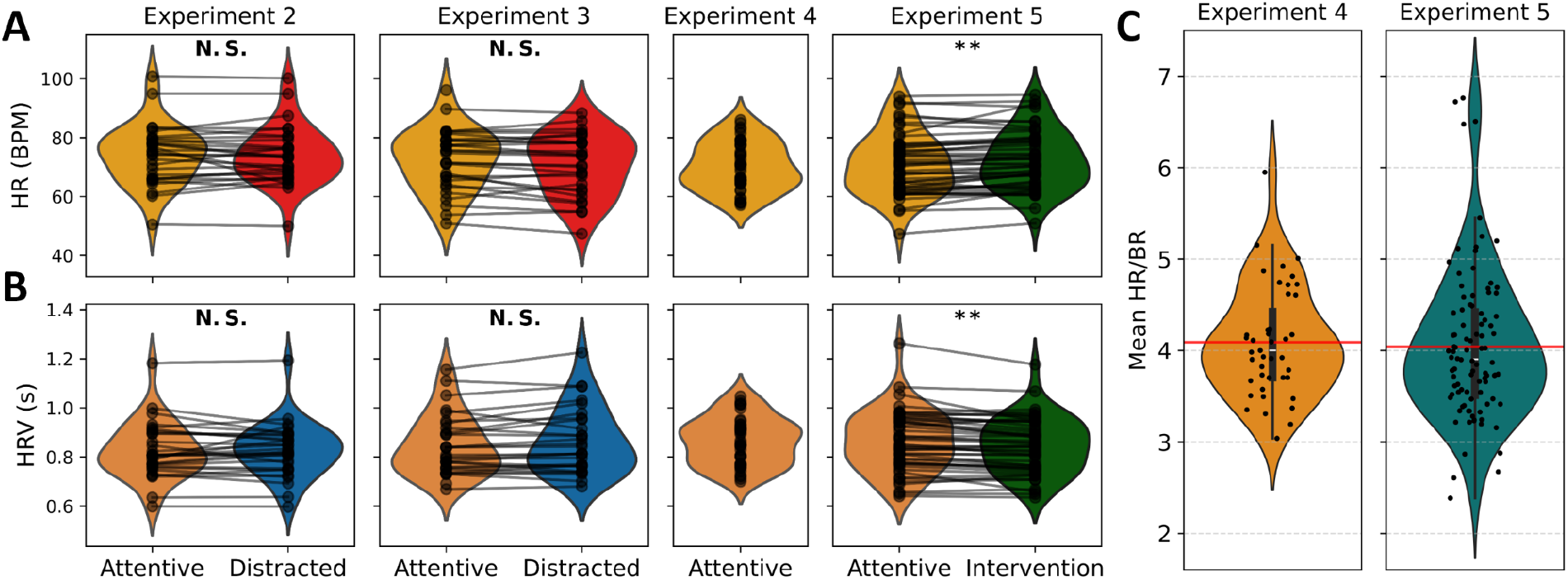
Distribution and changes in. **(A)** Heart rate and **(B)** Heart rate variability measured as the RMSSD (root mean square of successive differences) of R-R intervals across each video. Each line follows the respective subject in both experimental conditions watching the same video material in the two conditions. Measures were averaged across videos for each experiment/condition. For experiment 4 there was only one unique condition. **(C)** Average heart rate to breathing rate ratio for experiments that have respiration data (Exp 4, 5). (*) indicates a significant difference between the two groups at p<=0.05; (**) indicates the same at p<=0.01, and (N.S.) indicates no significant difference.

### Respiration Rate

We derive breathing rate from the respiration signal. The ratio of heart rate and breathing rate, known as Pulse-Rate Quotient (PRQ) has values consistent with previous literature^35^ (Fig. 6C, Exp 4: 3.77 ± 0.74, Exp 5: 3.73 ± 0.81, mean ± SD across subjects).

### Gaze, saccades, blinks and fixations

To validate the eye-tracker’s blink, fixation and saccade detection algorithm we investigate if these events align with the gaze position data. We observe that all interpolated periods (i.e. missing data) coincide with blinks. We also see that saccades coincide with large and rapid changes in gaze position, as expected^36^. In contrast, fixation durations are notably longer than those of saccades or blinks. During fixation periods, gaze position remains largely constant, as expected. The distinct patterns linked with each type of eye movement give us confidence in the effective functionality of the eye tracker, ensuring that it provides technically valid data for further analysis.

Blink rate increases when subjects are watching videos in the distracted and intervention condition compared to when subjects are attentive (Fig. 7A, Exp 1: t(26) = 2.09, p = 0.047, d’ = 0.41; Exp 2: t(30) = 2.58, p = 0.015, d’ = 0.47; Exp 3: t(27) = 2.04, p = 0.051, d’ = 0.39; Exp 5: t(47) = 0.27, p = 0.789, d’ = 0.04, within-subject paired t-test), while saccade rate is significantly reduced in the distracted and intervention conditions (Fig. 7B, Exp 1: t(26) = −7.28, p = 9.89·10^−8^, d’ = −1.43; Exp 2: t(30) = −8.79, p = 8.33·10^−10^, d’ = −1.61; Exp 3: t(27) = −5.73, p = 4.31·10^−6^, d’ = −1.10; Exp 5: t(47) = −3.46, p = 1.16·10^−3^, d’ = −0.50, within-subject paired t-test). These results are consistent with those of previous studies where an increase in cognitive load (distraction/intervention) is associated with the increase in the blink parameters such as the blink rate^37^, and a decrease in the saccadic movements - thus enabling higher focus^38^.

**Figure 7:**
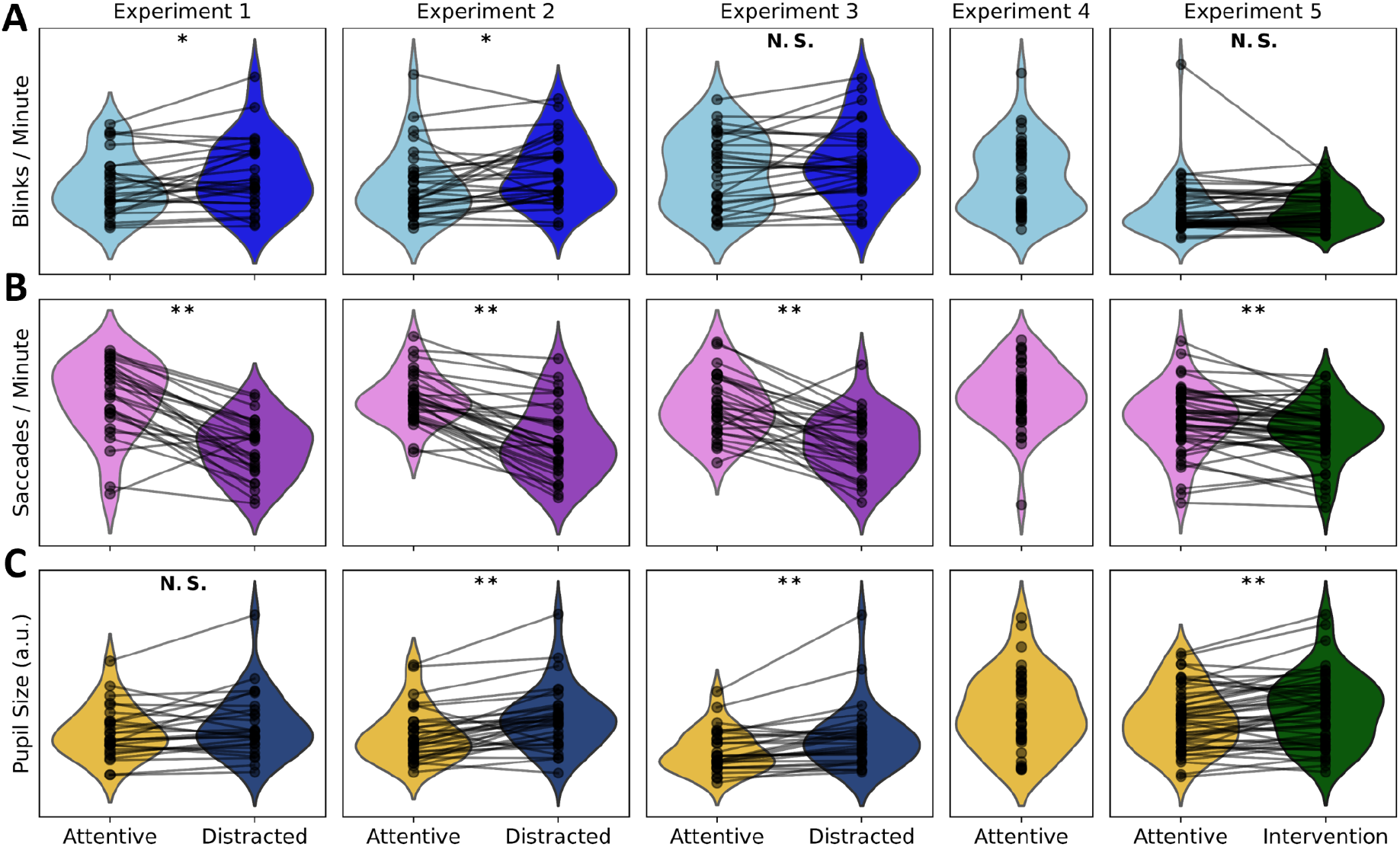
Distribution and changes in. **(A)** Blink Rate **(B)** Saccade Rate and **(C)** IQR (interquartile range) of Pupil Size. (*) indicates a significant difference between the two groups at p<=0.05; (**) indicates the same at p<=0.01, and (N.S.) indicates no significant difference.

### Pupil Size

Pupil size is higher in the distracted/intervention conditions (Fig. 7C), and this behaviour is consistent with existing literature that suggests a higher pupil area is associated with greater cognitive load^39,40^ and arousal^41^ (Fig. 7C, Experiment 1: t(26) = 1.82, p = 0.081, d’ = 0.36; Experiment 2: t(30) = 4.57, p = 7.8·10^−5^, d’ = 0.83; Experiment 3: t(27) = 3.81, p = 7.4·10^−4^, d’ = 0.73; Experiment 5: t(47) = 3.57, p = 8.4·10^−4^, d’ = 0.52, within-subject paired t-test).

To summarise, the following within-subject factors showed significant changes in a paired t-test across experimental conditions:

**Heart rate increased** in the intervention condition in Experiment 5 (p = 1.9·10^−4^, d’ = 0.59).

**Heart Rate Variability decreased** in the intervention condition in Experiment 5 (p = 6.6·10^−5^, d’ = −0.64).

**Blink rate increased** in the distracted/intervention conditions in Experiments 1,and 2 (Exp 1: p = 0.047, d’ = 0.41; Exp 2: p = 0.015, d’ = 0.47).

**Saccade rate decreased** in the distracted/intervention conditions in Experiments 1, 2, 3, and 5 (Exp 1: p = 9.89·10^−8^, d’ = −1.43; Exp 2: p = 8.33·10^−10^, d’ = −1.61; Exp 3: p = 4.31·10^−6^, d’ = −1.10; Exp 5: p = 1.16·10^−3^, d’ = −0.50).

**Pupil size increased** in the distracted/intervention conditions in Experiments 2, 3, and 5 (Exp 2: p = 7.8·10^−5^, d’ = 0.83; Exp 3: p = 7.4·10^−4^, d’ = 0.73; Exp 5: p = 8.4·10^−4^, d’ = 0.52).

## Usage Notes

EEG data files are stored in .bdf format and can be accessed using pyEDFlib^42^ or mne-bids^43^ libraries on Python, or EEGLAB^44^ on MATLAB. Physiological and behavioral signals are stored as compressed .tsv (tab-separated value) files, and can be accessed using the pandas^45^ library on Python. All metadata files are stored in .json, and the scores for digit-span, ASRS and stimuli quizzes are stored as .tsv files in the /phenotype directory of each experiment.

We encourage researchers to explore and utilize this dataset for advancing knowledge in cognitive and physiological sciences. This multimodal physiological signal dataset has potential to offer insights in various research areas

## Code Availability

We have released the code to reproduce the figures shown in this manuscript in the following github repository: https://github.com/madjens/bbbd-figures. We also released the code to download the datasets using either Python or MATLAB, along with scripts to visualize data, which can be found in the Tutorials section of the BBBD website: https://bbbd.pythonanywhere.com/tutorials-page.

## Acknowledgements

We acknowledge the National Science Foundation Grant DRL-1660548 for supporting this project, and also thank the many individuals that participated in these experiments.

## Author Contributions

J.M and L.C.P designed the experiments. J.M collected the data. J.M analyzed the data. N.K.K transformed the data into BIDS, technically validated it, and built the website that hosts the dataset. J.M N.K.K and L.C.P wrote the manuscript.

